# VR HMD color calibration and accurate control of emitted light using Three.js

**DOI:** 10.1101/2024.08.02.606304

**Authors:** Killian Duay, Yoko Mizokami, Takehiro Nagai

## Abstract

Virtual Reality (VR) can be used to design and create new types of psychophysical experiments. Its main advantage is that it frees us from the physical limitations of real-life experiments and the hardware and software limitations of experiments running on 2D displays and computer graphics. However, color calibration of the displays is often required in vision science studies. Recent studies have shown that a standard color calibration of a Head-Mounted Display (HMD) can be very challenging and comes with significant drawbacks. In this paper, we introduce a new approach that allows for successful color calibration of an HMD and overcomes the disadvantages associated with other solutions. We utilize a new VR engine, Three.js, which offers several advantages. This paper details our setup and methodology, and provides all the elements required to reproduce the method, including the source code. We also apply our method to evaluate and compare three different HMDs: HTC Vive Pro Eye, Meta Quest Pro, and Meta Quest 3. The results show that the HTC Vive Pro Eye performs excellently, the Meta Quest Pro performs well, and the Meta Quest 3 performs poorly.

## 1 Introduction

Virtual Reality (VR) allows researchers to design, create, and run experiments innovatively, opening new horizons and addressing some issues encountered in more traditional experiments. Indeed, it can be impossible or laborious to control certain experimental conditions in real-life experiments. For instance, changing the configuration of an experimental room or its lighting can be time-consuming and may potentially disrupt the experiment. Moreover, certain experimental conditions, such as introducing a mismatch between the light field of the background and the target object (Motoyoshi & Matoba, 2012), are impossible to produce in the real world. The utilization of computer graphics displayed on 2D screens has become a common approach to circumvent these limitations. However, this strategy introduces additional constraints, including a non-immersive environment and a limited field of view, which hinder the accurate replication of real-life conditions. VR makes it possible to overcome these physical limitations and offers an immersive environment that reproduces certain aspects of reality, such as a wide field of view and depth perception.

Color calibration refers to the procedure of establishing a relationship between monitor input (*RGB* reflectances) and monitor output (emitted light) and vice versa (Brainard, 1989). One of the purposes of such calibration is to present a desired luminance and chromaticity on the display. It is often required in experiments. Recent studies have shown the complexity of performing a standard calibration and accurately controlling the colors on a Head-Mounted Display (HMD) (Díaz-Barrancas, Gil-Rodríguez, Aizenman, Bayer, & Gegenfurtner, 2023; Gil Rodríguez et al., 2022; Toscani et al., 2019; Zaman, Sarker, & Tavakkoli, 2023). They explained that the underlying causes are attributable to both hardware and software factors. On the hardware side, HMDs are composed of near-eye displays (NED) and complex optical elements that make them difficult to measure with a spectroradiometer or a colorimeter (Gil Rodríguez et al., 2022). On the software side, VR experiences are executed through 3D engines relying on complex rendering pipelines involving post-processing routines that break the conditions necessary for a successful standard calibration (Gil Rodríguez et al., 2022; Toscani et al., 2019). As explained by Toscani et al. (2019), these conditions are the luminance additivity, chromaticity constancy, and channel constancy of the displays. These display characteristics are the criteria to be respected for standard calibration, as explained in older studies (Brainard, 1989; Brainard, Pelli, & Robson, 2002). Toscani et al. (2019) showed that the main software issue comes from the tone mapping process. Tone mapping is a post-processing routine used to approximate the appearance of High Dynamic Range (HDR) images on Standard Dynamic Range (SDR) displays. Although tone mapping improves the realism of a 3D scene, this disrupts the luminance additivity, chromaticity constancy, and channel constancy of the displays, making standard calibration unsuccessful. While VR offers significant advantages to researchers, the inability to calibrate an HMD poses substantial challenges in fields such as vision research, which necessitate precise control over the physical colors emitted by a display.

Nevertheless, the previous studies not only raised problems but also proposed solutions to meet the necessary conditions for calibration. Gil Rodríguez et al. (2022) demonstrated how to use a spectroradiometer equipped with a special VR lens in order to effectively measure an HMD and overcome the hardware issue. Díaz-Barrancas et al. (2023) showed how to calibrate an HMD by disabling tone mapping in Unreal Engine or Unity, addressing the software problem. Toscani et al. (2019) also demonstrated how to calibrate an HMD by disabling tone mapping in Unreal Engine. However, even though disabling tone mapping seems like the right solution, Zaman et al. (2023) explained that disabling it globally has a disadvantage. Specifically, it makes the overall rendering appear unrealistic as it changes the behavior of all shaders, materials, and objects in the scene. They suggested that it would be, therefore, beneficial to be able to selectively control which objects in a scene should undergo tone mapping and which do not. By doing so, we could accurately control stimulus colors while maintaining a realistic global scene and background. They proposed a solution and demonstrated how to calibrate an HMD using Unreal Engine and a custom double pathway in the rendering pipeline. This solution allows the tone mapping of the global VR scene but not of specific objects such as the stimulus in a VR experiment. However, although this approach is very interesting and addresses the previously mentioned problem, it has two disadvantages in our view. Firstly, the authors use unlit materials to implement the solution. Unlit materials are self-emissive and thus unaffected by the illuminant in the scene. Their color always remains the same regardless of the illuminant. This can be a significant drawback in experiments such as color constancy studies, where the influence of the illuminant on color perception is investigated. Secondly, this approach seems very complex and difficult to reproduce.

In this paper, we introduce and present a new approach that overcomes all the previously mentioned problems. We demonstrate that it is possible to perform a successful color calibration of an HMD using standard materials (as opposed to unlit materials) while the tone mapping is still applied to the rest of the scene. The strength of this solution lies in what we call modular tone mapping: we can easily choose which objects in a 3D scene should be affected by tone mapping and which should not. This modular tone mapping is an accessible and easy setting that can be implemented in two lines of code, as demonstrated in the code that we provide. This approach allows us to overcome all the previously mentioned problems. Firstly, it enables a global scene to appear realistic under tone mapping. Secondly, it provides accurate control over the physical color of objects of interest (such as the stimulus in an experiment) thanks to successful calibration. Finally, it ensures that all objects in the scene are adequatly influenced by the illuminant and changes in the illuminant (including the stimulus in an experiment). Additionally, our solution offers the possibility to globally disable tone mapping, as was done in the previously cited studies. Our solution is therefore available in two options: option *A* with tone mapping globally disabled and option *B* with tone mapping locally disabled. Each of these options has its own advantages and disadvantages, implying different use case scenarios. We describe the flexibility of our solution in more detail in the following sections. Furthermore, this solution has the advantage of being easily reproducible, and the source code and instructions are provided. It can be accessed at https://gitlab.com/kikdu/paperhmd-calibration-scene.

Moreover, unlike other studies, our solution employs Three.js instead of more conventional platforms such as Unreal Engine or Unity. Three.js is a JavaScript application programming interface (API) built on top of OpenGL, WebGL, and WebXR. It allows access to the graphics processing unit (GPU) and runs 3D and VR applications in a web browser. Thanks to the performance of the V8 JavaScript engine and browsers built on Chromium, Three.js is able to render applications with very high graphics quality and a high frame rate, similar to desktop applications. In addition to providing a solution using a different tool and bringing more options and diversity to the research community, we believe that Three.js has other advantages compared to Unreal Engine or Unity. Firstly, the source code is readily intelligible, allowing full control over everything that happens in the scene. This is attributable to the low-level nature of the library, which closely interfaces with the OpenGL API. In contrast, Unity and Unreal Engine can sometimes appear as black boxes where precise control over all internal processes is not achievable. Secondly, Three.js VR experiences run in a web browser, making them highly portable to any execution environment thanks to their device-independent nature. This can be advantageous in specific scenarios, such as in a non-tethered experiment where the use of an HMD allows for independence from a computer. Finally, although it does not have a graphics editor, Three.js can be less complex to set up and master than Unity or Unreal Engine, which can sometimes feel overwhelming for non-experts.

In the following sections, we describe our setup and methodology in detail to facilitate easy reproduction of our procedure. We explain how we measured the HMDs, characterized the displays, performed the actual calibration, and validated it. Similar to recent studies (Díaz-Barrancas et al., 2023; Gil Rodríguez et al., 2022; Toscani et al., 2019; Zaman et al., 2023), we demonstrated our approach through two successive and distinct setups. Firstly, we demonstrated and illustrated the problem of global tone mapping. Secondly, we introduced and applied our solution, showing how it overcomes this and other previously mentioned problems. We validated our solution by performing the same calibration test as done in Gil Rodríguez et al. (2022) and Toscani et al. (2019), ensuring consistency and comparability with other recent studies. Additionally, we applied our method to evaluate and compare three different HMDs: HTC Vive Pro Eye, Meta Quest Pro, and Meta Quest 3. While the first is common, and recent studies have shown that it can be calibrated (Gil Rodríguez et al., 2022; Toscani et al., 2019; Zaman et al., 2023), the latter two HMDs have not yet been assessed in research. Therefore, it is important to evaluate whether they can be calibrated and utilized in experiments that require precise color control. The results showed that the HTC Vive Pro Eye performed excellently, the Meta Quest Pro well, and the Meta Quest 3 poorly. We also compared the color reproduction fidelity across the entire lens of each HMD. We demonstrated that while we can precisely control the color in the sweet spot of the lenses, it is much more challenging to do so in the peripheral areas. To our knowledge, this is the first study to characterize and compare color reproduction across the entire lens of HMDs. In our view, this is a crucial detail to consider in VR psychophysical experiments involving colors and a significant flaw in other related works.

## 2 Methods

### 2.1 Calibration of a 3D environment

Calibrating a 3D environment differs from a standard calibration, as defined by Brainard (1989). A standard calibration procedure is based on the relationship between *RGB* values in each pixel and the characteristics of the display to represent a desired color on the display. On the other hand, calibrating a 3D environment relies on a “relative” color space composed of *RGB* reflectance values of objects, in which the emitted colors depend not only on the display characteristics but also on scene lighting and the spatial arrangement of objects. As explained by Toscani et al. (2019), the color of the light, its intensity, and the distance between the light and the calibration target define the relative color space (luminance range and gamut) of the 3D environment. Figure 1 illustrates this explanation: all planes share identical *RGB* reflectances ([1, 1, 1]^*T*^); however, their color on the display varies with their distance from the light source (denoted by the white point) in Figure 1a and Figure 1c. Additionally, changes in the intensity of the light source affect the colors as well, as depicted in Figure 1a and Figure 1b. Thus, in this case, the aim of color calibration is to establish a relationship between *RGB* reflectances (input) and the light reflected by an object in the 3D scene (i.e., the output light presented on the display). Because this output also depends on the position of the object in space and the characteristics of the lighting, calibrating a 3D environment is done for a specific area and layout of that space. Additionally, if these characteristics are changed after the calibration, for example, if the target patch is moved or the light source is modified, the calibration is no longer valid. Consequently, if the calibration is done for a research experiment, it is important to adapt the spatial layout of the calibration room to match the spatial layout of the virtual 3D room hosting the experiment. The code we provide allows this adaptation easily.

**Figure 1:**
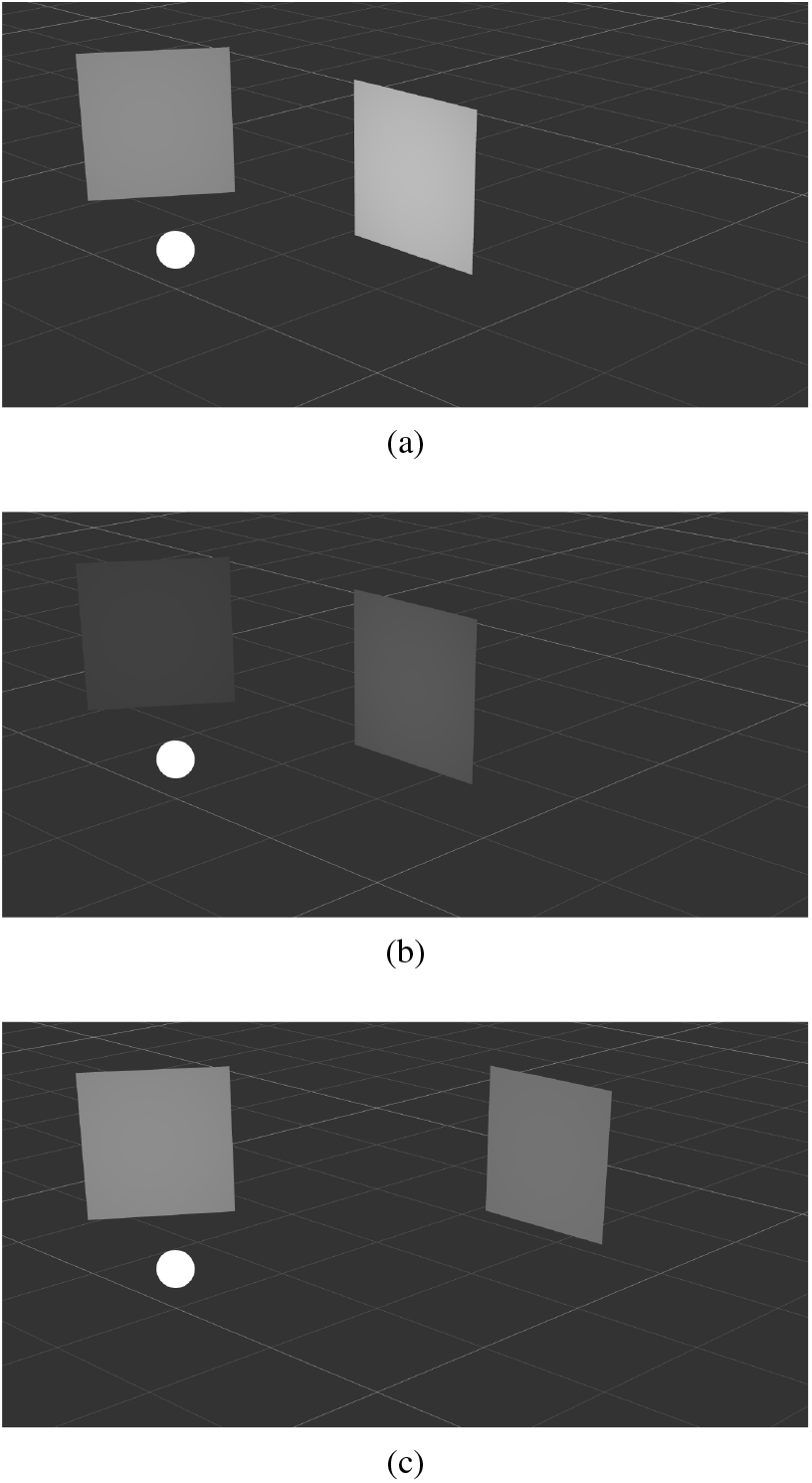
Examples of 3D scenes with different spatial layouts and light intensities. In (a), the light intensity is 5 candelas (cd). In (b), the light intensity is 1 cd. In (c), the light intensity is 5 cd. The white point represents the light source position. All planes have the same *RGB* reflectances [1, 1, 1]^*T*^.

By using the procedure explained here, one can only calibrate a specific object in the scene. After calibration, it is possible to determine the object’s physical color in advance, but this is not possible for all other objects and pixels in the scene. This is because they are located at different distances from the light source and might not share the same properties. Moreover, as explained in Section 1, the calibration of a 3D VR scene typically occurs in game engines such as Unity or Unreal Engine. Toscani et al. (2019) and Gil Rodríguez et al. (2022) showed that it is necessary to disable the tone mapping post-processing in those engines in order for the display to achieve sufficiently constant levels of luminance additivity, chromaticity constancy, and channel constancy for a standard calibration. Although this can be used to calibrate an HMD, Zaman et al. (2023) claimed that this solution is not ideal, as this makes the 3D scene look unrealistic and unnatural. Although they proposed a solution allowing tone mapping to be disabled on specific objects only, their implementation is highly complicated and uses unlit materials. In contrast, our solution aims to address both of these issues.

Finally, another important consideration when employing VR for research experiments is that the engines utilized in our approach or in the previously cited studies compute reflected *RGB* colors solely through linear *RGB* calculations. They do not perform physically correct computations based on spectral distributions. Gil Rodríguez et al. (2024) explain that *RGB* reflected light is calculated using the following formula:

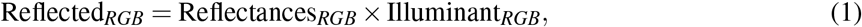

where “Reflectances” is defined as the proportion of light of each channel that an object reflects and is a property of the object regardless of illumination. While this rendering technique is employed by the majority of game engines and is capable of generating environments that closely replicate reality with high visual fidelity, it is crucial to clarify that it does not constitute an accurate simulation of real-world color computation based on light spectrum. This limitation is primarily due to the necessity for these engines to render in real-time and maintain high frame rates, which, as of the current writing, cannot be achieved by physically correct simulation engines.

### 2.2 Our method

Our solution using Three.js can be used in two different ways:

A. Tone mapping globally disabled, similar to Toscani et al. (2019) and Gil Rodríguez et al. (2022),
B. Tone mapping locally disabled, similar to Zaman et al. (2023) but using standard lit materials.

These two alternatives can be interchanged by adjusting a simple parameter. In both options, the outcome is the disabling of tone mapping for the target object undergoing calibration, thereby enabling a successful color calibration process. However, they each have different advantages and disadvantages. In option *A*, tone mapping is globally disabled for the entire scene. This ensures consistent object colors throughout the scene, meaning luminance and color there can be physically correct. However, the scene has a narrow luminance dynamic range, making it look unrealistic or unnatural if the scene is in a high dynamic range in nature. This option should be used for experiments requiring strict and consistent color behavior, such as color perception experiments. In option *B*, tone mapping is applied to the entire scene but locally disabled for specific objects. This allows for more natural-looking 3D scenes with a higher range of luminance due to the tone mapping still being applied to them. The disadvantage of this solution is that tone mapping is not applied uniformly throughout the scene; therefore, two objects with the same reflectances but different tone mappings will have different colors. This might change the way observers perceive the scene and the relative colors. This option can be used for experiments where it is important to calibrate the color of specific objects while keeping the rest of the scene looking natural and realistic.

In addition, unlike Zaman et al. (2023), our solution uses standard lit materials in both options, meaning that the target objects being calibrated still react to the illuminant or changes in the illuminant.

Figure 2 shows an example of the visual impact of both options on a simple scene. In Figure 2a, the scene appears saturated due to the absence of tone mapping and the larger dynamic range of physical luminance. In Figure 2b, the scene appears more realistic and natural thanks to the tone mapping globally enabled, but a standard calibration cannot be performed successfully because of this setting. In Figure 2c, tone mapping is globally enabled but locally disabled for the second helmet (from the left). As a result, the scene looks more natural, and the second helmet can still be calibrated since it is not affected by tone mapping. Please note that the scenes shown in Figure 2 are used only to illustrate our tone mapping settings and do not represent the scene used for the actual calibration presented in this study.

**Figure 2:**
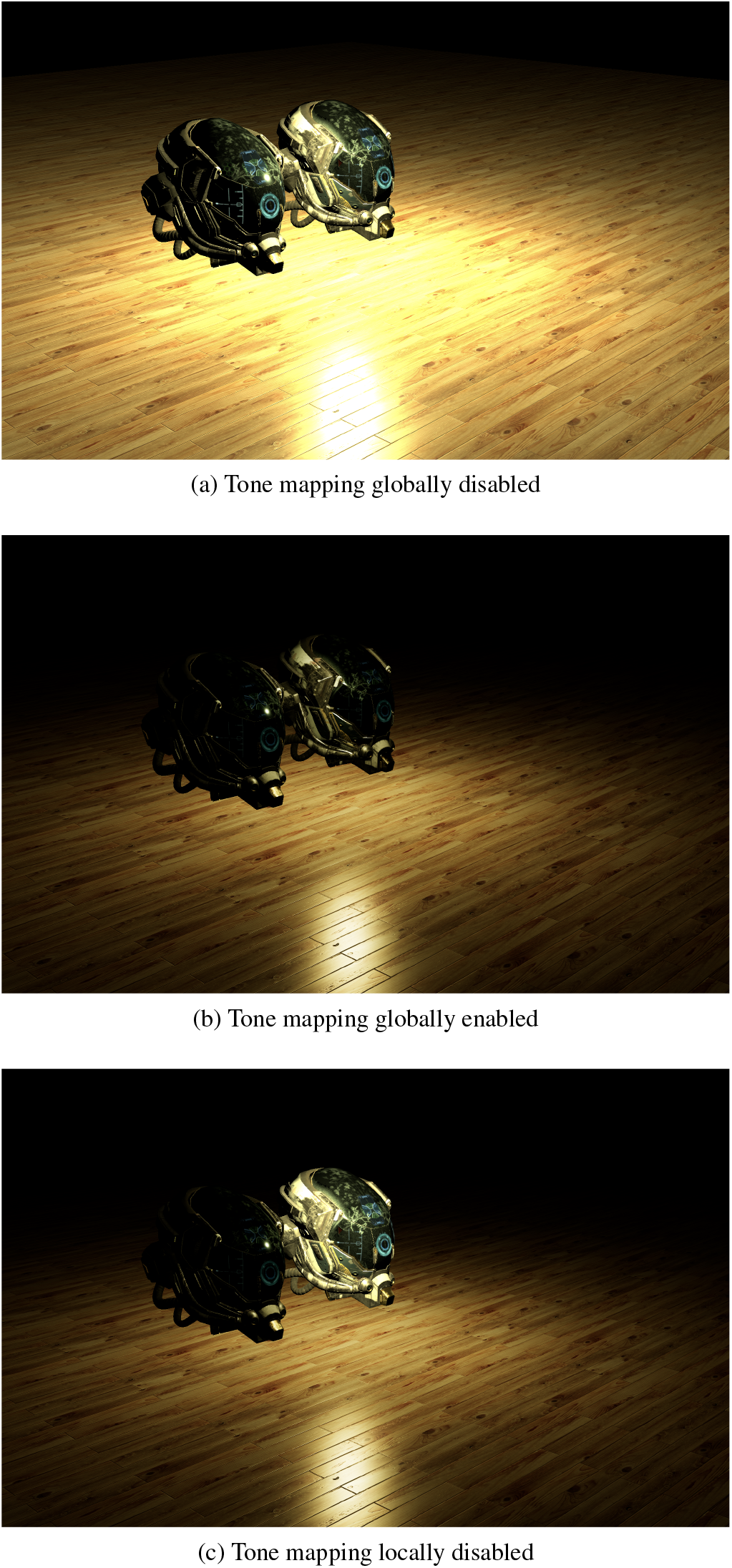
Examples of 3D scenes with different tone mapping settings. In (a), tone mapping is globally disabled. In (b), tone mapping is globally enabled. In (c), tone mapping is globally enabled for the entire scene but locally disabled for the second helmet (from the left).

The methodology of the calibration process is the same for both options. Similar to other studies (Gil Rodríguez et al., 2022; Toscani et al., 2019; Zaman et al., 2023), our approach consists of four steps. First, we take the measurements of the HMD using a spectroradiometer. Second, we characterize the display and its luminance additivity, channel constancy, and chromaticity constancy based on those measurements. Third, if the conditions are met in the second step, we process the data and compute the calibration. Finally, we validate the calibration through a test. Each of these steps and the supporting setup are detailed in the following sections.

### 2.3 Hardware and software

We utilized and compared three different HMDs: HTC Vive Pro Eye, Meta Quest Pro, and Meta Quest 3. Table 1 summarizes some of the technical specifications of these HMDs. The VR program was executed on a desktop computer (Windows 10, Intel(R) Core(TM) i9-10900K CPU @ 3.70 GHz, 32 GB of RAM, Nvidia Quadro RTX 4000 8GB). The HTC Vive Pro Eye was tethered to the PC via its original connector and the SteamVR desktop application. The Meta Quest Pro and Meta Quest 3 are standalone HMDs, but in this study, we tethered them to the PC using a Meta Quest Link and the Meta Quest desktop application. For the VR software and 3D engine, we developed our solution with Three.js (release r159). As previously explained, Three.js can be used to develop and execute VR experiences in a web browser. The web browser we used in our procedure was Google Chrome.

**Table 1:**
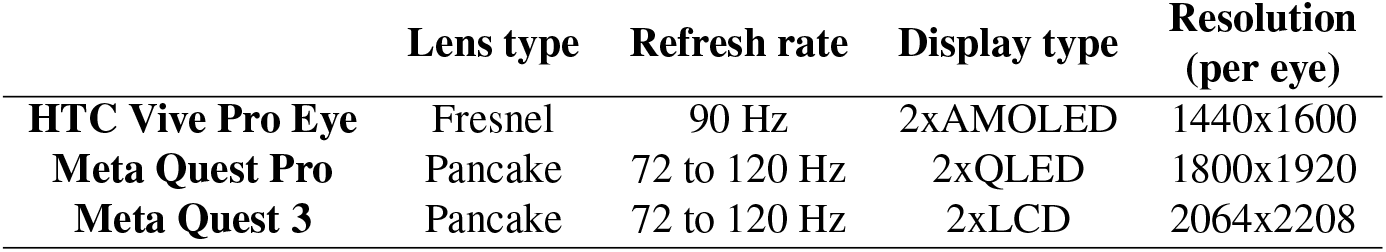
Technical specifications of the HMDs used in this study.

|  | Lens type | Refresh rate | Display type | Resolution<br>(per eye) |
| --- | --- | --- | --- | --- |
| **HTC Vive Pro Eye** | Fresnel | 90 Hz | 2xAMOLED | 1440x1600 |
| **Meta Quest Pro** | Pancake | 72 to 120 Hz | 2xQLED | 1800x1920 |
| **Meta Quest 3** | Pancake | 72 to 120 Hz | 2xLCD | 2064x2208 |

To take the measurements, we utilized a Topcon SR-5000 HM spectroradiometer equipped with a specific macro lens and a retrofitted lens (LM5JC1M, Kowa Optronics), which allowed us to achieve a clear focus inside the HMD by placing it very close to the HMD lens. Figure 3 shows the measurement setup. The spectroradiometer was controlled through proprietary software. We positioned the spectroradiometer the same distance from the HMD lens as our eye would be when wearing the HMD (2 centimeters from the surface of the lenses). However, we could not replicate the same setup for the Meta Quest Pro. Due to its construction and headband, the lens of the spectroradiometer could approach the lens of the Meta Quest Pro only up to a maximum of 10 centimeters from the surface of the lenses, as shown in Figure 3b.

**Figure 3:**
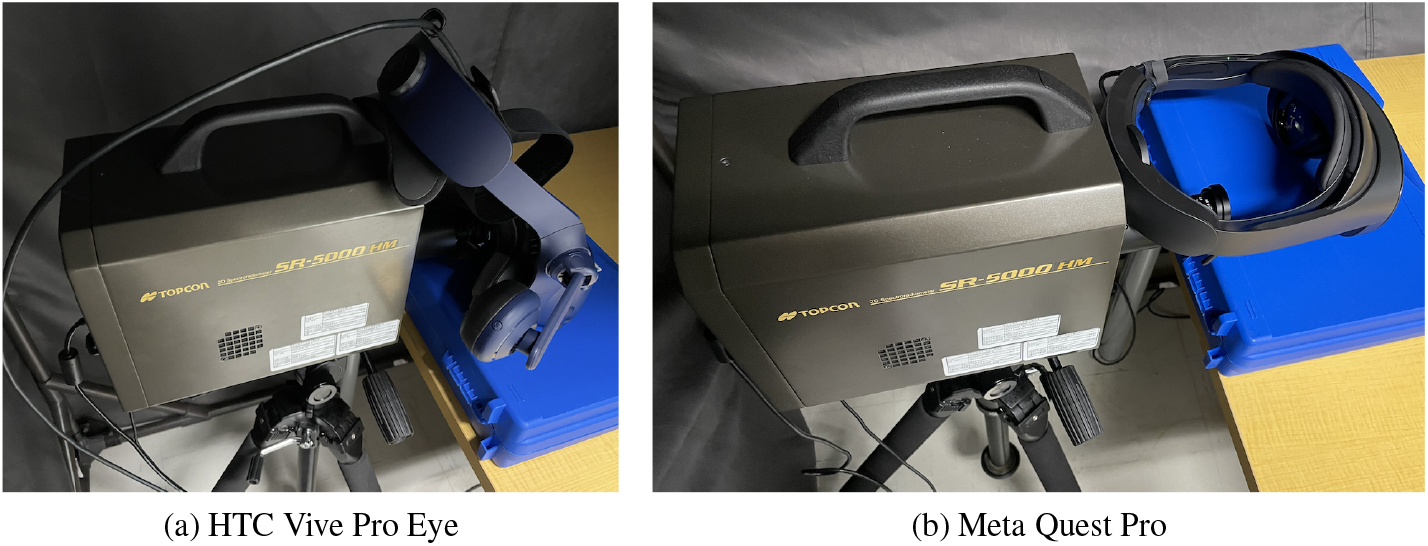
Setups of the spectroradiometer and the HMDs for measurements: (a) HTC Vive Pro Eye and (b) Meta Quest Pro.

Finally, we also used an external monitor connected to the desktop computer to control and orchestrate the entire setup. This complete setup included the VR program running inside the HMD and the spectroradiometer software. The measured HMD and the spectroradiometer were positioned in a darkroom, while the computer and monitor used to control the whole setup were positioned outside of the darkroom.

### 2.4 Calibration scene and measurements

We developed a 3D scene designed for HMD calibration. Figure 4 shows the scene. It is empty, with a background set to *RGB* reflectances [0, 0, 0]^*T*^, an ambient neutral light set to *RGB* intensities [1, 1, 1]^*T*^, a point light source set to *RGB* intensities [1, 1, 1]^*T*^, and a stimulus object. In a 3D environment that does not use ray-tracing rendering techniques and bouncing lights, the ambient light illuminates all objects in the scene equally, simulating indirect lighting and Global Illumination (GI). On the other hand, the point light source acts as a traditional direct lighting. The stimulus object is a flat square with a perfectly Lambertian material. It is located in front of the camera in the virtual world and aligned with the sweet spot of the right lens of the HMD (we used the right eye for our measurements). In VR, the notion of the camera corresponds to the position of the user’s eyes in the virtual world. The point light is located behind the camera and is aligned with the stimulus and the camera. The stimulus object and the point light follow the camera orientation, and they are always positioned respectively in front of and behind the camera, aligned with the sweet spot of the lens, regardless of the position or orientation of the HMD in the real world. This detail is important for achieving stable and consistent measurements. The sweet spot of the lens is where the user’s eye is located when wearing the HMD and is also where the focus and the luminance are highest.

**Figure 4:**
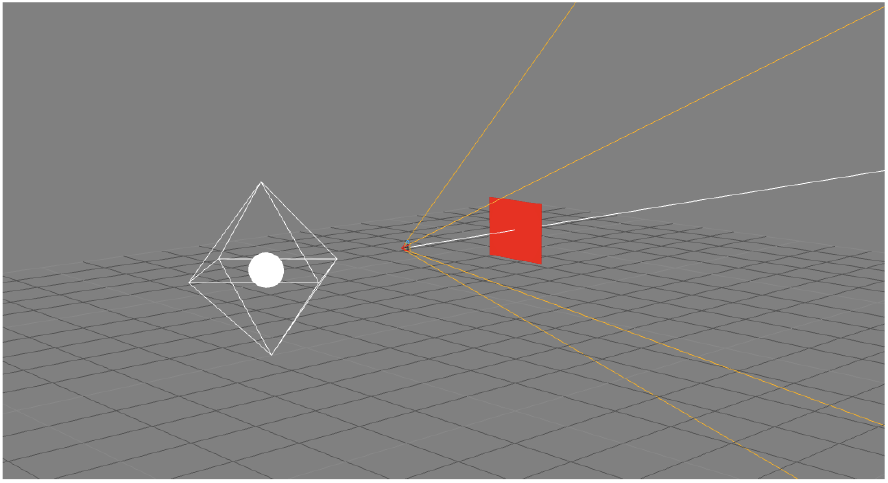
Calibration room. The stimulus is shown in red. The white point and white diamond represent the light position. The intersection of the orange and white lines indicates the VR camera position. The white line represents its orientation, and the orange lines represent its field of view. The grid, white diamond, white point, and orange and white lines are only visualization aids in editor mode. They are not present in the actual view displayed in the HMD.

We measured the focus and the luminance using the spectroradiometer. We observed that the sweet spot is perfectly in the middle of the lenses in the HTC Vive Pro Eye and slightly in the top-right and top-left areas of the lenses, respectively, for the left eye and right eye in the Meta Quest Pro and Meta Quest 3. This can be seen in the first row of Figure 14. Figure 5 shows an example of the view of the HTC Vive Pro Eye from the spectroradiometer and how we aligned the VR camera, the stimulus, the HMD lens, and the spectroradiometer for measurements. The displayed cross is the helper we used to align everything before measurements.

**Figure 5:**
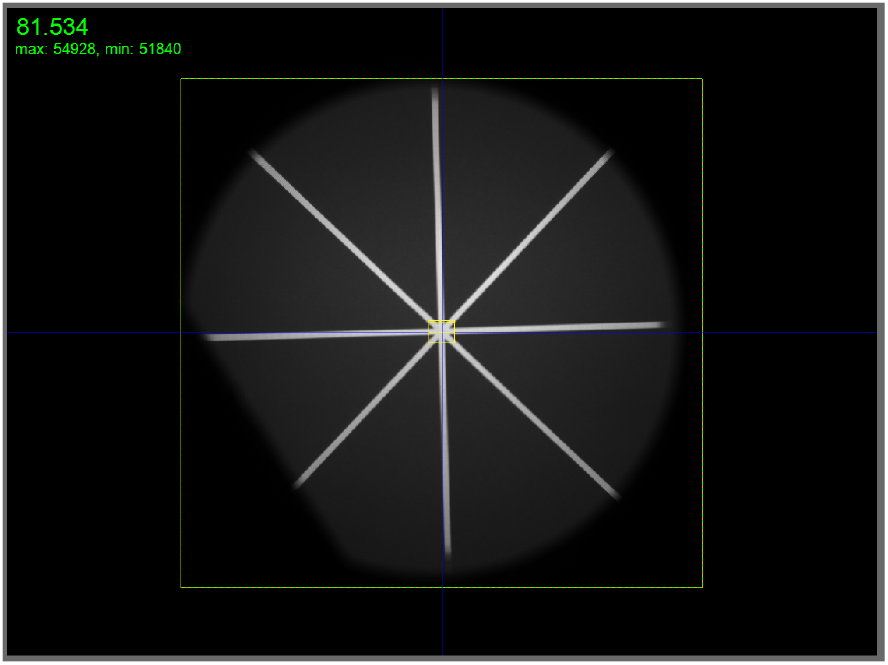
View of the stimulus captured by the spectroradiometer. The VR camera, stimulus, HMD lens, and spectroradiometer are perfectly aligned.

Using the arrow keys on the keyboard, it is possible to change the color of the stimulus. To take measurements for calibration, the stimulus displays the red channel from *RGB* reflectances [0, 0, 0]^*T*^ to [1, 0, 0]^*T*^ in 6 successive and incremental steps. After the red channel, it repeats the same process for the green and blue channels. Finally, it displays all three channels together from *RGB* reflectances [0, 0, 0]^*T*^ to [1, 1, 1]^*T*^ in 6 incremental steps. This calibration scene allows us to take 24 measurements. In Three.js, the *RGB* reflectances are given in linear RGB color space, but the scene is configured to use sRGB as the ouput color space. In combination with the gamma properties of the display, which are expected to follow the gamma properties defined in sRGB, the measured output luminance for each channel should increase linearly, and simple linear interpolation would provide an accurate prediction of luminance for any *RGB* reflectances. For this reason, we judged that six measurements per channel were sufficient.

### 2.5 Calibration process

Following the methodology outlined by Toscani et al. (2019), we first characterized the behavior of the software and the display to verify that they comply with standard calibration conditions as defined by Brainard (1989). We evaluated the luminance additivity, channel constancy, and chromaticity constancy of the red, green, and blue channels based on the measurements. We used the calibration scene in order to take the 24 measurements of the red, green, blue, and gray channels separately, as described in Section 2.4. The spectroradiometer allowed us to capture the chromaticity, luminance, and spectral distribution of each stimulus.

If the necessary calibration conditions were met, the calibration was then performed following a standard calibration method such as described by Koida (2021), Toscani et al. (2019) and Wilson and Hua (2021). We used the 24 measurements previously taken to compute the calibration matrix and functions. First, we performed a linear interpolation of the luminance measurements and built look-up tables (LUT) that map *RGB* reflectances to luminance outputs and vice versa. Before proceeding with the interpolation, we subtracted the luminance value measured with the *RGB* reflectances [0, 0, 0]^*T*^ from each other luminance measurement. We performed that operation and built a look-up table for each channel separately, such as:

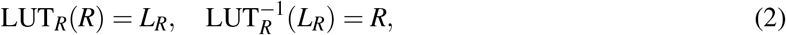

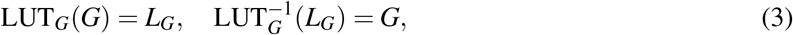

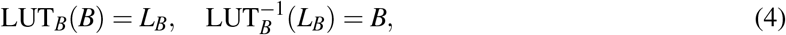

where *R, G* and *B* are the *RGB* reflectances of primary colors, and *L*_*R*_, *L*_*G*_ and *L*_*B*_ are the luminance ouput. Second, we computed the calibration matrix such as:

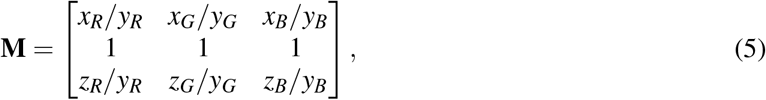

with (*x*_*R*_, *y*_*R*_, *z*_*R*_), (*x*_*G*_, *y*_*G*_, *z*_*G*_), and (*x*_*B*_, *y*_*B*_, *z*_*B*_) being the chromaticity coordinates of the displayed stimulus of *RGB* reflectances [1, 0, 0]^*T*^, [0, 1, 0]^*T*^, and [0, 0, 1]^*T*^, respectively. Finally, the calibration functions are given by:

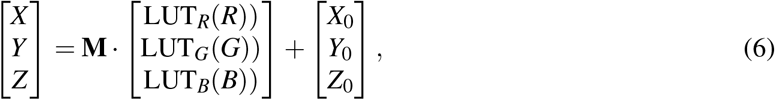

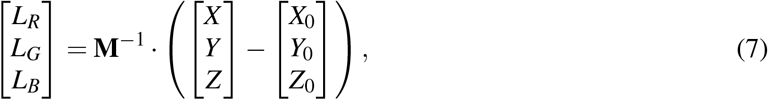

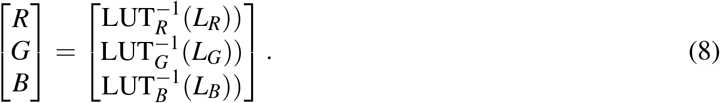

As a result, we can predict the XYZ tristimulus for any given *RGB* reflectances using Equation 6. We can also reciprocally compute the *RGB* reflectances needed to produce any XYZ tristimulus using Equation 7 and Equation 8. In the Equations 6 and 7, the vector [*X*_0_,*Y*_0_, *Z*_0_]^*T*^ is the measured XYZ tristimulus values for the *RGB* reflectances [0, 0, 0]^*T*^. Additionally, the relationship between the final RGB values of the image (not the reflectances) and XYZ tristimulus values can also be described in the same way as with standard calibration.

### 2.6 Validation scene and measurements

The validation scene is almost identical to the calibration scene. The only distinction lies in the set of stimulus colors. Instead of displaying the red, green, and blue channels separately, the validation scene displays a set of colors used for validating the calibration process. This set was inspired by Toscani et al. (2019). The *RGB* reflectances of the validation set represent a large cube and a small cube in the three-dimensional RGB color space. The large cube vertices range from *RGB* reflectances [0.2, 0.2, 0.2]^*T*^ to [0.8, 0.8, 0.8]^*T*^, and the small cube vertices range from *RGB* reflectances [0.4, 0.4, 0.4]^*T*^ to [0.6, 0.6, 0.6]^*T*^. The validation color stimulus set, therefore, consists of 16 different colors spanning the entire RGB gamut and luminance range.

### 2.7 Validation process

We used the same validation method as Gil Rodríguez et al. (2022) and Toscani et al. (2019). First, we used the validation scene to take the measurements of the 16 colors in the validation set, as described in Section 2.6. Then, we compared those measurements with the predictions of our calibration function introduced in Section 2.5. To compare them, we computed the color difference Δ*E*_00_ between the 16 measured values and the 16 predicted values by using CIEDE2000. These 16 color differences were then averaged into a single color difference score, referred to as the validation score. A validation score smaller than 2 represents an excellent color calibration since it is below the threshold of perceptually noticeable differences between two colors for the human visual system (Gil Rodríguez et al., 2022; Sharma & Bala, 2017).

### 2.8 Experimental setups

To demonstrate and illustrate our methodology, we applied it to two different experimental setups:

- Three.js with tone mapping globally enabled (for both the entire scene and the measured target object)
- Three.js with tone mapping globally enabled (for the entire scene) but locally disabled (for the measured target object)

The aim of the first setup is to again demonstrate and illustrate the problem of tone mapping exposed in Toscani et al. (2019), Gil Rodríguez et al. (2022) and Zaman et al. (2023). The aim of the second setup is to demonstrate that our approach allows for successful calibration. The second setup demonstrates both options *A* and *B*, introduced in Section 2.2, simultaneously. It is unnecessary to demonstrate the two options independently, as their outcomes would be identical. Indeed, as explained in Section 2.2, tone mapping is disabled for the target object of the calibration in both cases. Evaluating both options would be akin to testing the same parameters twice in succession, yielding identical results. Therefore, the differences between options *A* and *B* cannot be demonstrated quantitatively, but they can be appreciated qualitatively in Figure 2.

## 3 Results

### 3.1 Calibration with tone mapping globally enabled

The first experimental setup aims to demonstrate the issue of tone mapping. In this setup, the tone mapping was simply enabled globally for the entire scene and every object in the scene. As the aim is to demonstrate the issue of color calibration at the software level (i.e. with tone mapping globally enabled, in contrast and comparison with the next section where it is disabled), we used only one HMD as an example (HTC Vive Pro Eye).

#### 3.1.1 Characterization of the display

Following the process described in Section 2.5, we initially characterized the display and the software. Figure 6 shows the relationship between *RGB* reflectances and luminance output for the three channels separately (red, blue, and green). In addition, the sum of the RGB luminance (i.e., gray) is also shown in the rightmost panel; the dashed line represents the luminance additivity (sum of luminance independently measured for red, green, and blue channels), and the bold line shows the measured luminance when all RGB channels were presented simultaneously. The three primary channels are not increasing linearly, which would have been the expected behavior, as explained in Section 2.4. Ideally, with excellent additivity behavior, the dashed line would perfectly align with the solid line in the rightmost panel. However, the luminance additivity is found to be very poor in this setup. Figure 7 shows the spectra of the three channels individually for varying *RGB* reflectances. The continuous lines represent the measured spectra for the different *RGB* reflectances, while the dashed lines represent the spectra predictions obtained by multiplying the lowest measured spectrum by a linear scaling factor. Linear scalings of the lowest spectrum result in spectra that perfectly fit the spectra measured for higher *RGB* reflectances for the red and blue channels. This behavior suggests very good channel constancy for those channels. However, the green channel seems much less constant. Two extra peaks of radiance that are not linearly increasing can be observed. Finally, Figure 8 shows the chromaticity coordinates (*x, y*) of the three channels with normalized *RGB* reflectances higher than 0.20 on the CIE 1931 chromaticity diagram. In the case of good chromaticity constancy, the dots of the same color should overlap. The chromaticity constancy is satisfactory for the red and blue channels but poor for the green channel, as expected from the spectrum in Figure 7.

**Figure 6:**
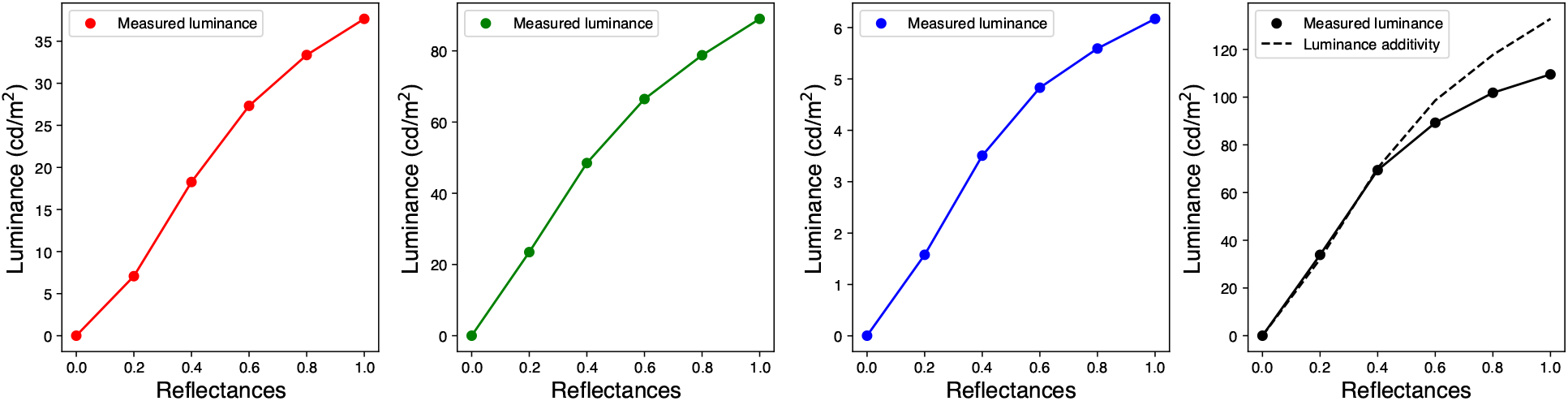
Relationship between *RGB* reflectances and luminance output for the three channels (red, blue, and green channels) separately with globally enabled tone mapping. Additionally, the rightmost panel shows the same for gray, where all *RGB* channels are simultaneously used (the continuous line) in addition to the sum of the luminance of red, green, and blue channels (the dashed line) to check luminance additivity.

**Figure 7:**
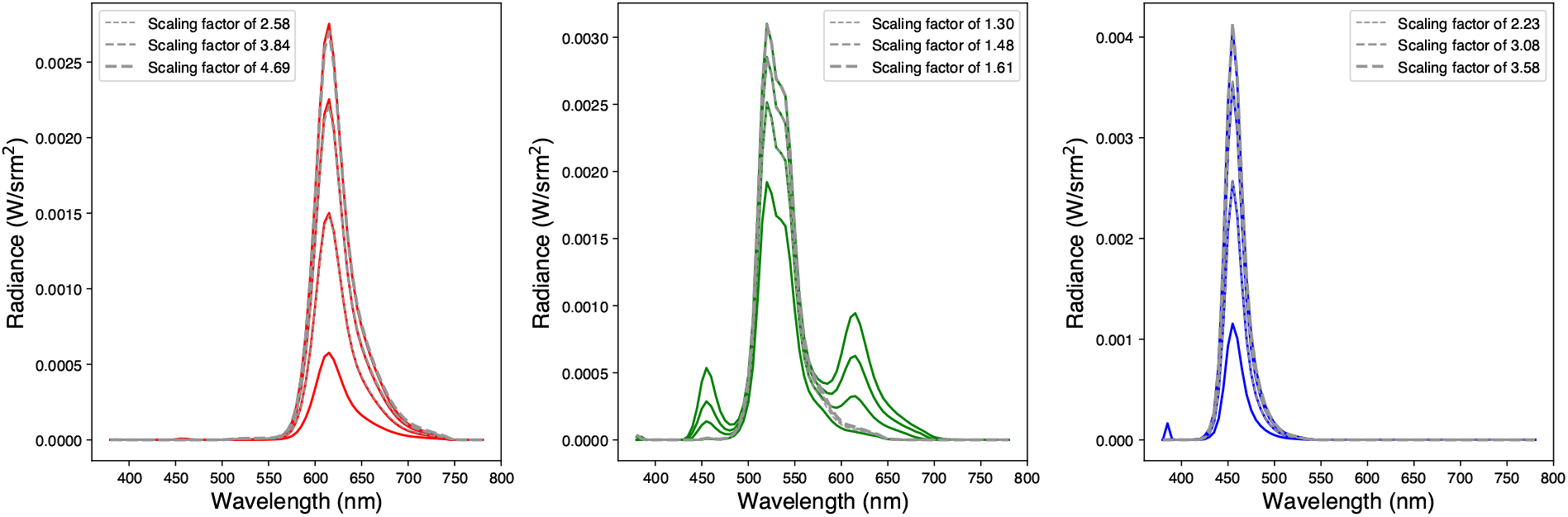
Measured spectra of three channels separately for different *RGB* reflectances (0.4, 0.6, 0.8, 1.0) with globally enabled tone mapping. The continuous lines represent the measured spectra for the different *RGB* reflectances. The dashed gray lines represent the predicted spectra obtained by linearly scaling the lowest spectrum by a factor specified in the legends.

**Figure 8:**
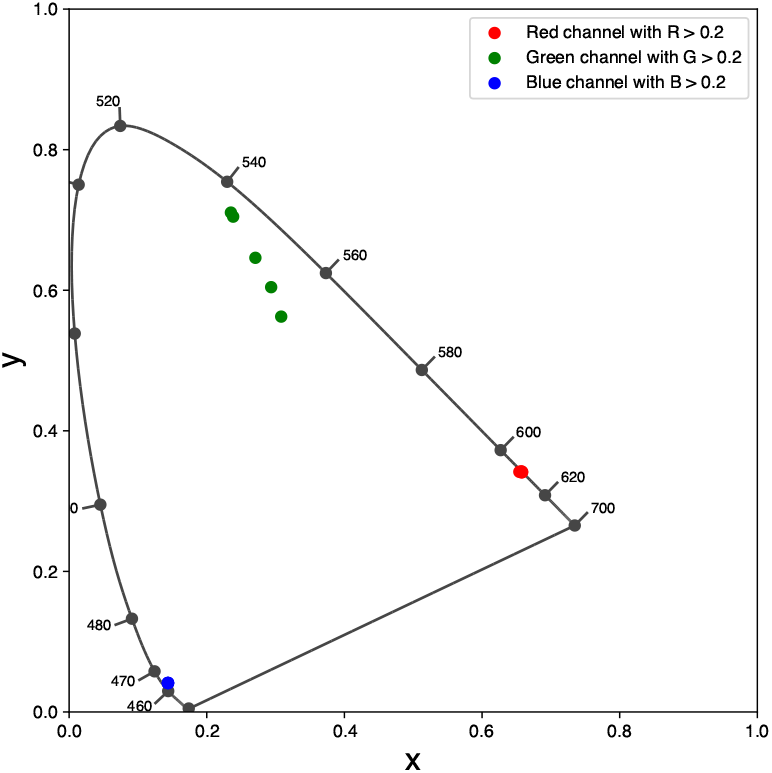
Chromaticity coordinates (x,y) of the three channels with *RGB* reflectances higher than 0.20 on CIE 1931 chromaticity diagram with tone mapping globally enabled.

#### 3.1.2 Calibration results and validation

This setup with tone mapping globally enabled showed poor luminance additivity, channel constancy, and chromaticity constancy. For those reasons, standard calibration is not expected to be performed effectively and not to yield satisfactory results. Nevertheless, we proceeded with the calibration to demonstrate this issue. We employed the calibration method described in Section 2.5 and the validation method described in Section 2.7. The outcome was an average Δ*E*_00_ of 14.9 as a validation score, indicating poor calibration performance and a significant, perceptible color difference between predictions and measurements. Figure 9 shows the discrepancies between predictions and measurements on the CIE 1976 UCS chromaticity diagram. The red crosses represent the measurements, and the black dots represent the predictions. It is evident that predictions and measurements do not closely align, which would be the expected result in a successful calibration.

**Figure 9:**
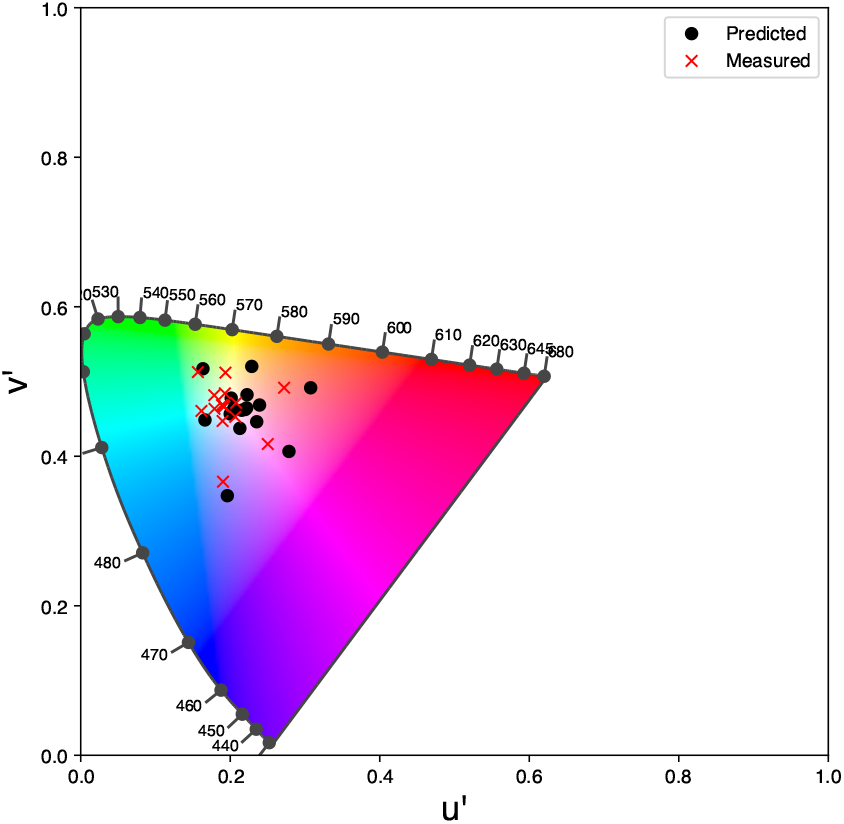
Chromaticity coordinates of 16 validation colors on CIE 1976 UCS chromaticity diagram with tone mapping globally enabled. The red crosses indicate the measured values, and the black dots indicate the values predicted by our calibration function.

### 3.2 Calibration with tone mapping locally disabled

The second experimental setup with tone mapping locally disabled is designed to illustrate and validate our approach and its advantages. In this setup, we measured and calibrated the three different HMDs described in Section 2.3.

#### 3.2.1 Characterization of the display

Following again the procedure described in Section 2.5, we first characterized the displays and the software. In this instance, Figure 10 demonstrates a very good linear relationship between *RGB* reflectances and luminance output for the HTC Vive Pro Eye, aligning with the expected behavior as previously discussed. The linearity is slightly less consistent for the blue channel of the Meta Quest Pro and the three primary channels of the Meta Quest 3. It is also noted that the Meta Quest 3 seems to reach a maximum level of luminance, which is not problematic and has been similarly observed by Gil Rodríguez et al. (2022) in a different setup. Regarding luminance additivity, the HTC Vive Pro Eye and the Meta Quest 3 exhibit slight over-additivity in high luminance, but this remains acceptable. On the other hand, the Meta Quest Pro shows slight sub-additivity, yet this is also still satisfactory. Figure 11 shows an excellent channel constancy for the HTC Vive Pro Eye, with similar results for the Meta Quest Pro. Although there are small extra peaks of radiance, they follow the linear scaling. However, the Meta Quest 3 shows much less channel constancy, with larger extra peaks of radiance that do not increase linearly. This suggests a lack of independent scaling for the primary channels, which may interfere with calibration. Finally, Figure 12 illustrates excellent chromaticity constancy across all three HMDs, with the dots nearly perfectly overlapping.

**Figure 10:**
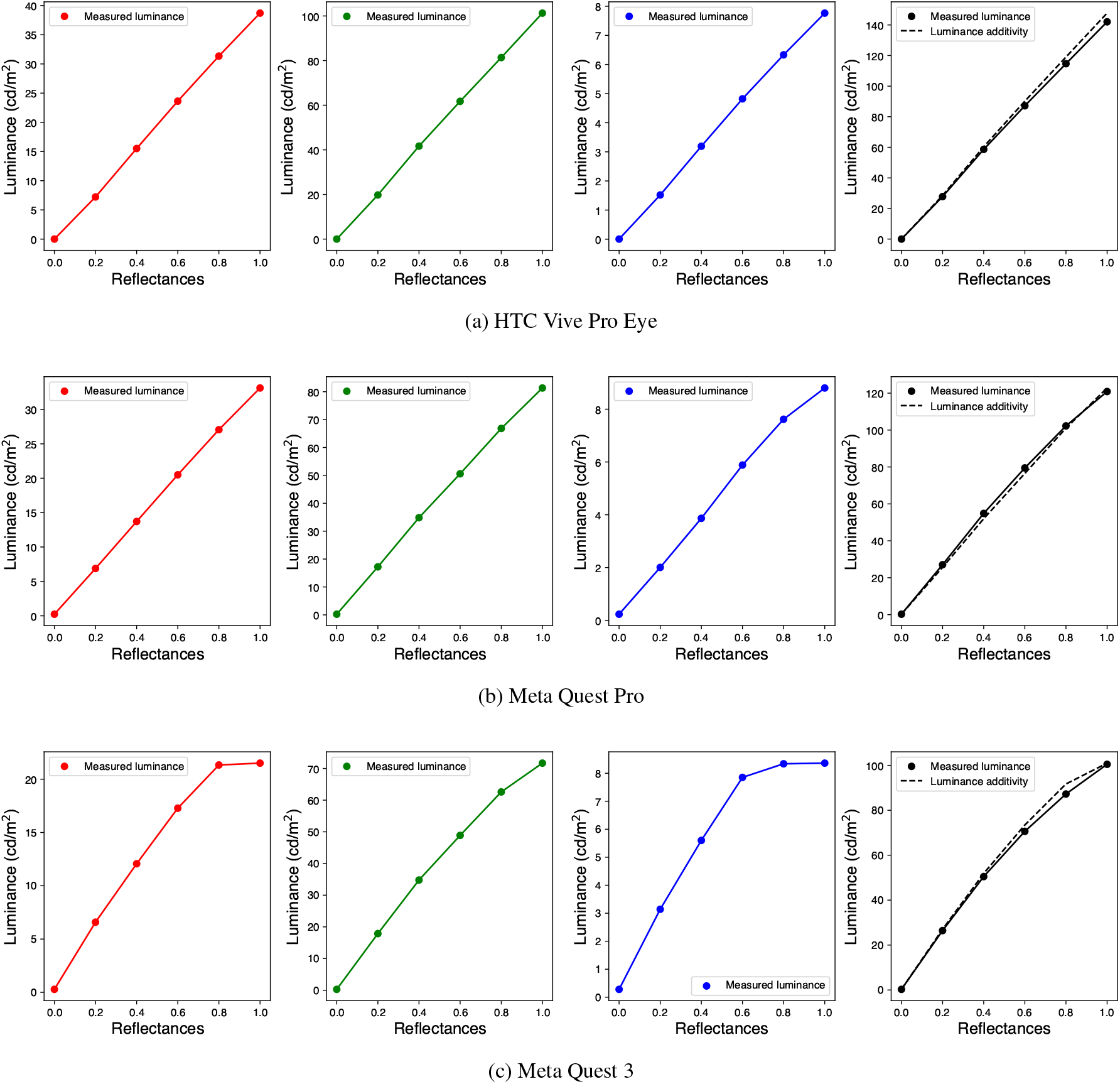
Relationship between *RGB* reflectances and luminance output for the three channels (red, blue, and green channels) separately with tone mapping locally disabled. The results for HTC Vive Pro Eye, Meta Quest Pro, and Meta Quest 3 are presented in (a), (b), and (c), respectively. Additionally, the rightmost panels show the same for gray, where all RGB channels are simultaneously used (the continuous line) and the sum of the luminance of red, green, and blue channels (the dashed line) to check luminance additivity.

**Figure 11:**
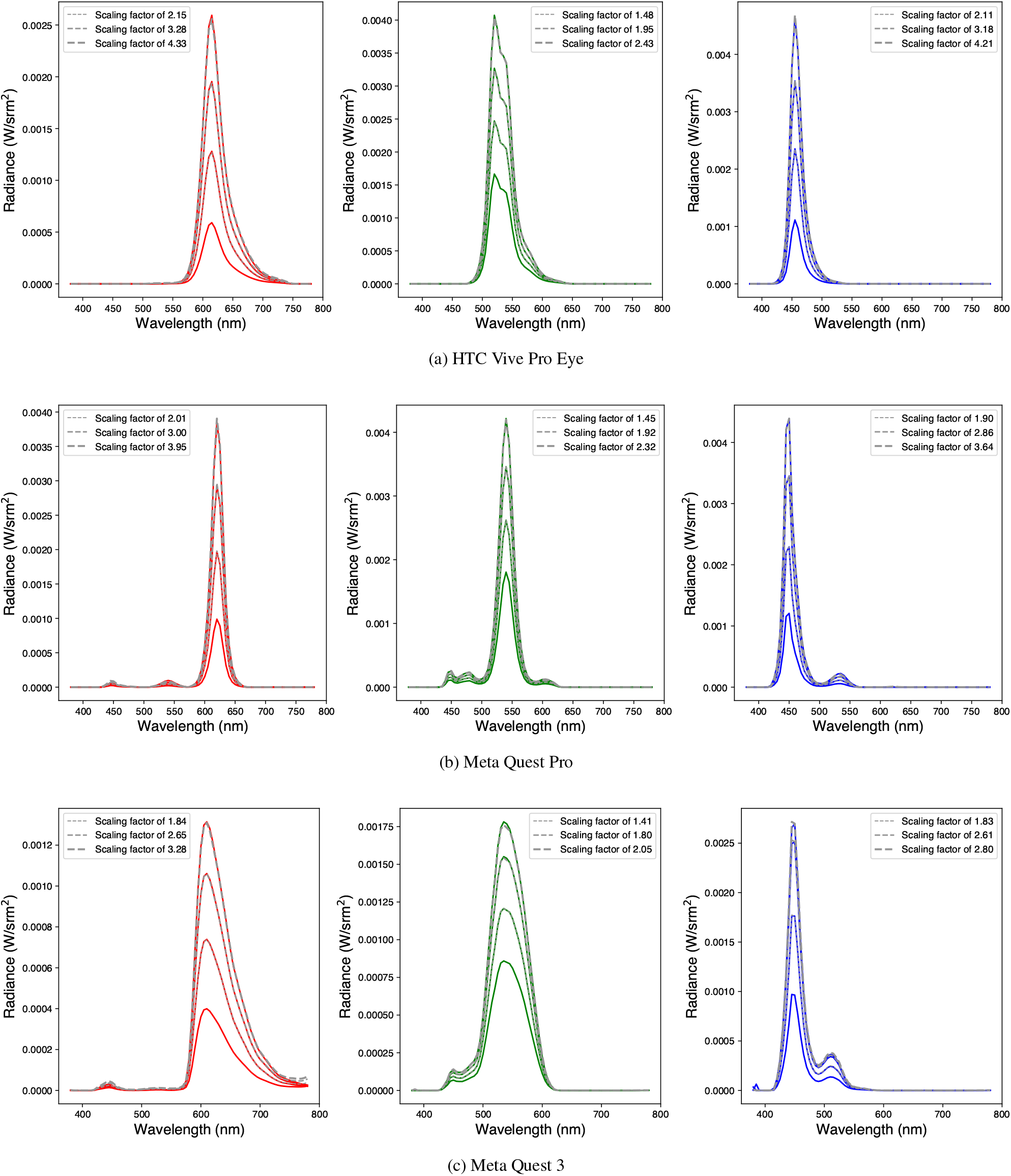
Measured spectra of each of the three channels for varying *RGB* reflectances (0.4, 0.6, 0.8, 1.0) with tone mapping locally disabled. The continuous lines represent the measured spectra for the different *RGB* reflectances. The dashed gray lines represent the predicted spectra obtained by linearly scaling the lowest spectrum by a factor specified in the legends. The results for HTC Vive Pro Eye, Meta Quest Pro, and Meta Quest 3 are presented in (a), (b), and (c), respectively.

**Figure 12:**
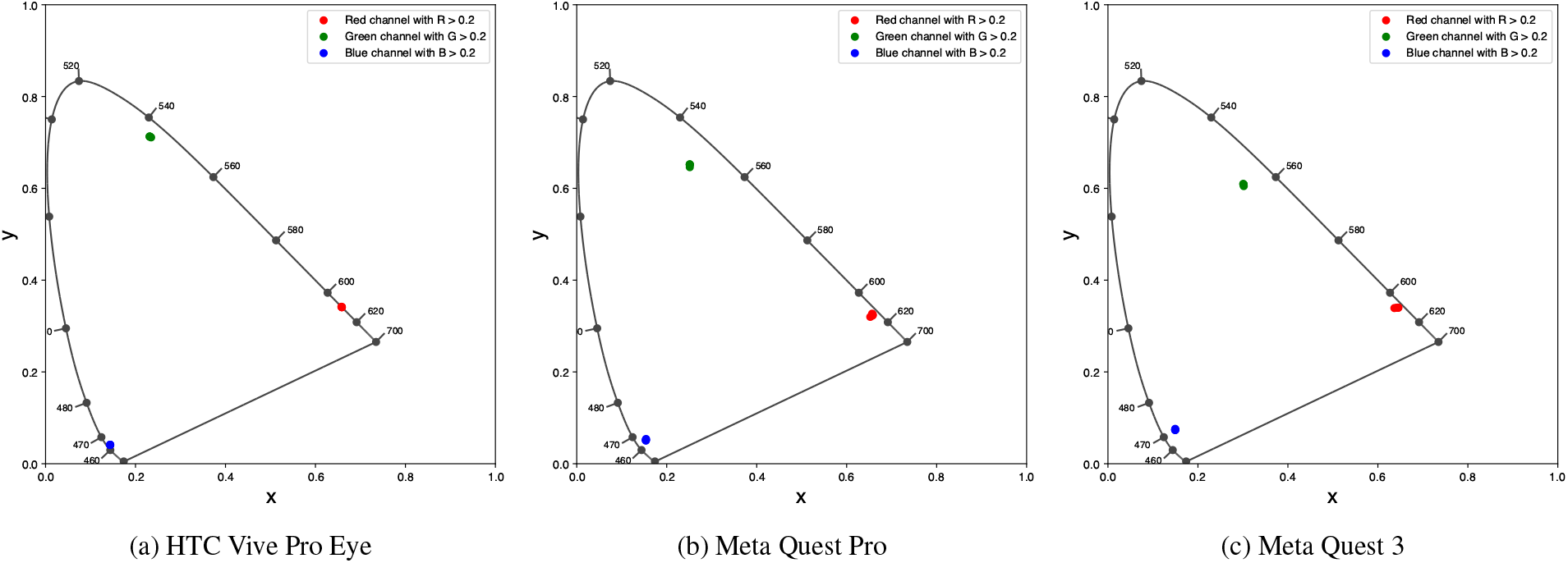
Chromaticity coordinates (x,y) of three channels with *RGB* reflectances higher than 0.20 on CIE 1931 chromaticity diagram with tone mapping locally disabled. The results for HTC Vive Pro Eye, Meta Quest Pro, and Meta Quest 3 are presented in (a), (b), and (c), respectively.

#### 3.2.2 Calibration results and validation

This characterization demonstrated satisfactory levels of luminance additivity, channel constancy, and chromaticity constancy for the three HMDs. With the conditions for standard calibration met, we subsequently applied the calibration procedure described in Section 2.5. To evaluate the results of the calibration, we ran the validation procedure described in Section 2.7. Figure 13 shows the results for all three HMDs on the CIE 1976 UCS chromaticity diagram. A perfect overlap between the red crosses and black dots on the HTC Vive Pro Eye diagram suggests that the calibration is successful and that the HMD and the software can produce and control colors effectively. Although slightly worse, the Meta Quest Pro also shows satisfactory results. On the other hand, the results for the Meta Quest 3 suggest that the calibration is not successful and is approximative. Indeed, the red crosses and black dots do not overlap in some areas of the diagram, such as the green or blue regions. Ultimately, we computed the validation score (Δ*E*_00_) as defined in Section 2.7. Table 2 shows the results for all three HMDs. These results confirm the suggestions made from the CIE 1976 UCS diagram. The HTC Vive Pro Eye performs excellently with a score of 1.0, which is below the threshold of perceptually noticeable difference for the human visual system (Gil Rodríguez et al., 2022; Sharma & Bala, 2017). The Meta Quest Pro performs less well, with a score of 3.9, a bit higher than the discrimination threshold. Finally, the Meta Quest 3 has an inferior score of 6.6.

**Table 2:** Mean average color differences Δ*E*_00_ between measured data and prediction data for HTC Vive Pro Eye, Meta Quest Pro, and Meta Quest 3 with tone mapping locally disabled.

| | Mean Average $\Delta E_{00}$ |
| --- | --- |
| HTC Vive Pro Eye | 1.0 |
| Meta Quest Pro | 3.9 |
| Meta Quest 3 | 6.6 |

**Figure 13:**
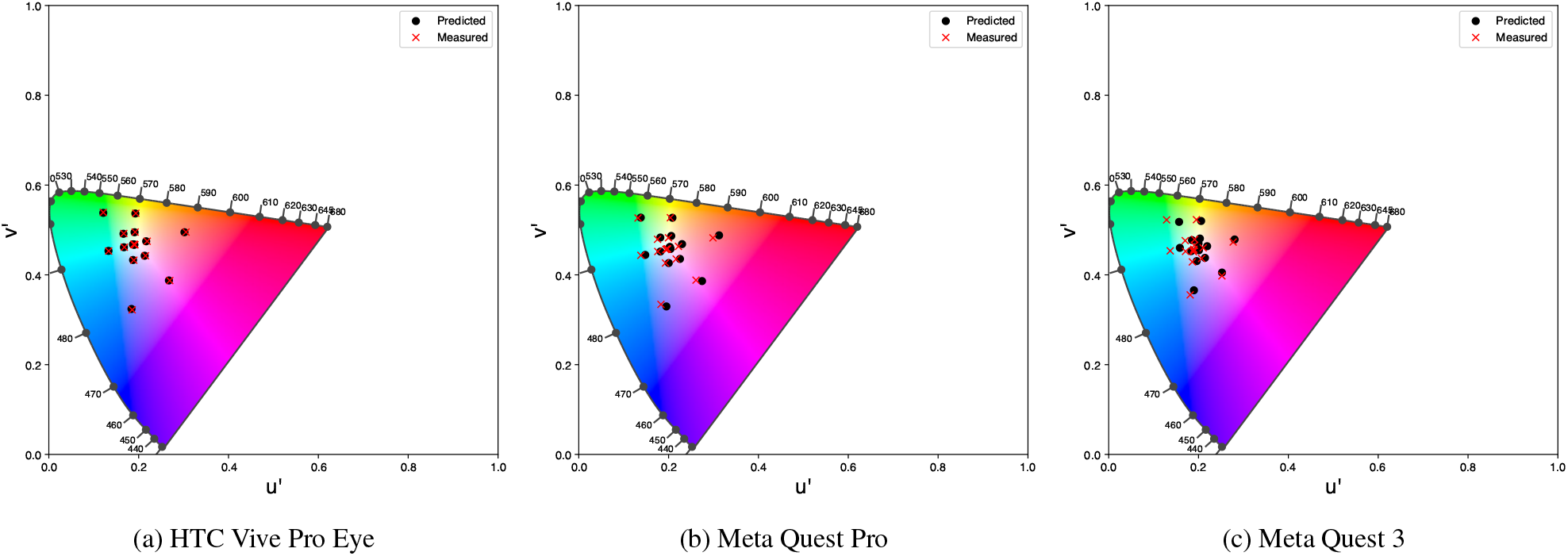
Chromaticity coordinates of 16 validation colors on CIE 1976 UCS chromaticity diagram with tone mapping locally disabled. The red crosses indicate the measured values, and the black dots indicate the values predicted by our calibration function. The results for HTC Vive Pro Eye, Meta Quest Pro, and Meta Quest 3 are presented in (a), (b), and (c), respectively.

These scores were obtained in the sweet spot of the lenses. We previously defined the sweet spot of the lens as the area where the user’s eye is positioned by default and where the luminance can reach the highest values. We also computed the validation score at other spots on the different lenses to compare the color reproduction fidelity across the entire lens area. Figure 14 illustrates these different scores for all three HMDs. It also shows the maximum luminance level of the lenses per area. As previously explained, we captured the data by placing the spectroradiometer at the same distance from the lens as our eye would be positioned if we were wearing the HMD (except for the Meta Quest Pro). We captured the maximum luminance levels by measuring a perfectly white stimulus covering the entire lens. It is observed that the best average score for each HMD is where the luminance reaches the highest value, which is also where the user’s eye is theoretically positioned. It is also observed that the color reproduction fidelity of the Meta Quest Pro and Meta Quest 3 varies drastically across the entire lens. They achieve scores exceeding 20, indicating that the colors in those regions are completely different. These results are directly explained by the large variation in luminance across the lenses, which is considered to be induced by lens distortion. On the other hand, the reproduction is more stable on the HTC Vive Pro Eye, and the differences are smaller. Although smaller, the differences in color reproduction fidelity across the three HMDs are still significant. These results suggest that it is much more challenging to control colors in the peripheral regions, especially in the Meta Quest Pro and Meta Quest 3, even if it is possible to precisely control the color at the center of the lenses.

**Figure 14:**
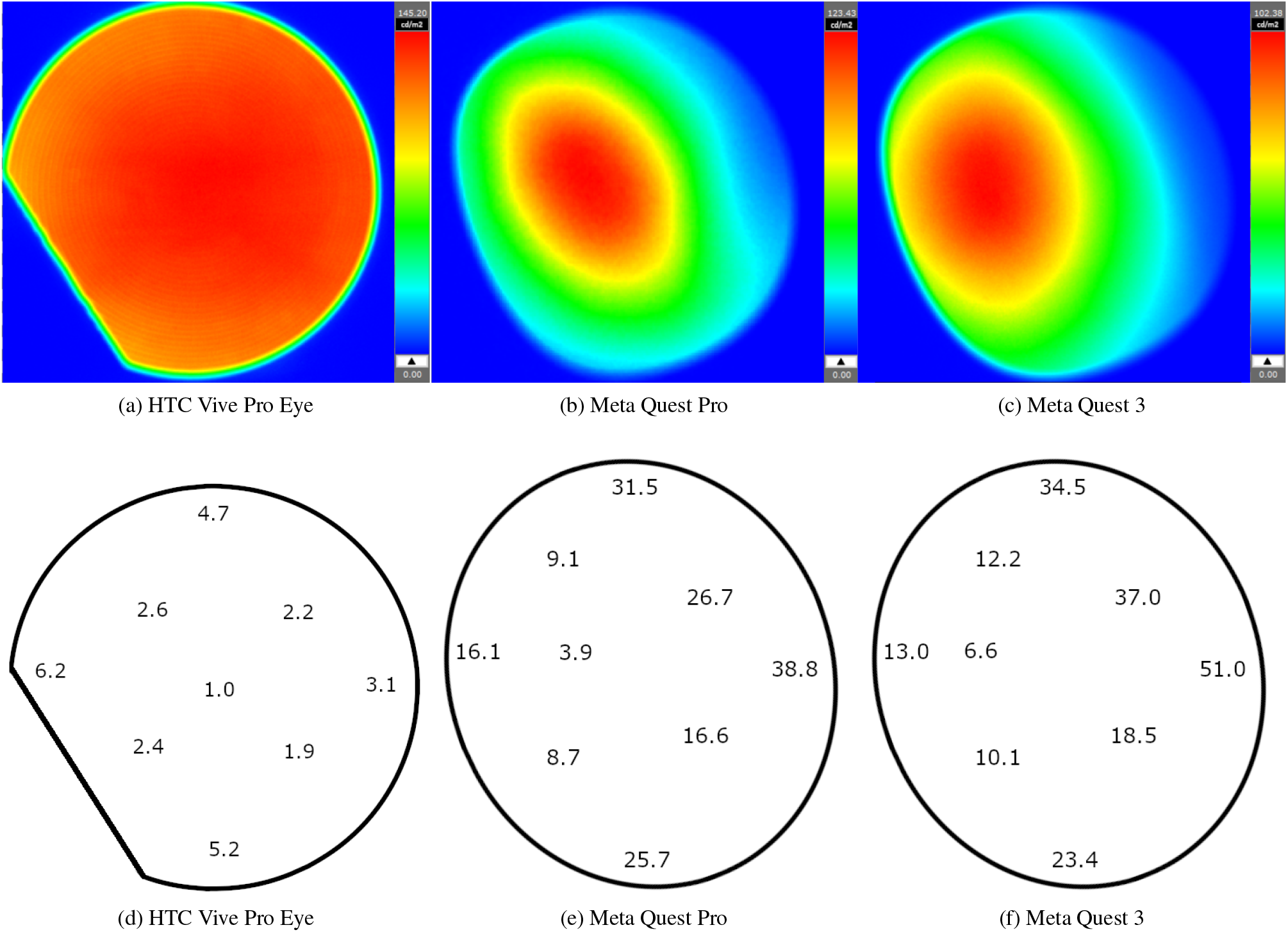
Color uniformity. First row: luminance across the area of the lens. Every scale belongs to the image on the left of it, and the unit is *cd/m*^2^. Second row: mean average color differences Δ*E*_00_ across the area of the lens with tone mapping locally disabled. The results for all three HMDs are shown in columns.

## 4 Discussion

This study aimed to propose a framework for HMD color calibration that overcomes some of the issues of previous solutions. We presented a comparative analysis between different HMDs and their outcomes under varying tone mapping conditions, highlighting the strengths of our approach. Our results offer insightful implications for HMD choices and calibration strategies.

The findings from the globally enabled tone mapping setup highlighted significant limitations in the HMD’s ability to maintain consistent color channel behavior. The poor luminance additivity, channel constancy, and chromaticity constancy underscore a fundamental problem in achieving reliable calibration under these conditions. The validation process yielded a high average Δ*E*_00_ of 14.9, indicating a noticeable discrepancy between expected and observed color outcomes. This substantial discrepancy suggests that global tone mapping introduces complexities that prevent traditional calibration methods from working. These results are consistent with previous studies (Gil Rodríguez et al., 2022; Toscani et al., 2019; Zaman et al., 2023).

Contrasting these results, the locally disabled tone mapping setup provided a more favorable environment for calibration. Here, the tone mapping applied globally except on the stimulus allowed for more controlled conditions. This technique demonstrated a marked improvement in calibration accuracy, as reflected by the near-perfect overlap between predicted and measured values and the average Δ*E*_00_ of 1.0 in devices like the HTC Vive Pro Eye. A calibration result of this nature may be considered ideal, as the average color difference falls below the perceptual threshold of the human visual system. These results highlight the potential of our approach and modular tone mapping in overcoming the limitations observed previously. Moreover, our solution also allows tone mapping to be globally disabled as Toscani et al. (2019). We have provided a qualitative comparison of options *A* and *B* that shows the visual difference of a scene between them (Figure 2). Option *A* should be used in scenarios where precise color control and consistency within the scene is important, such as in color constancy experiments. On the other hand, option *B* should be used in experiments where it is important to have precise control over the color of a target object while maintaining a very natural and realistic background. It should also be remembered that in both cases, only a small spatial portion of the scene can be calibrated, due to the nature and constraints of calibrating a 3D environment. However, as previously explained, our method and calibration functions also allow to predict luminance and chromaticity of any object in the scene based on the RGB values output by the display. Finally, it should also be noted that this method can accurately control the color of a flat patch only. If the object has a complex shape, it is only possible to precisely control the color of a restricted portion of its surface, while the remaining part of the object color will be altered by the shading or shadows.

Our analysis also revealed device-specific performance variations that merit consideration. The HTC Vive Pro Eye consistently showed excellent performance in terms of calibration and color reproduction accuracy. The Meta Quest Pro showed good results. In contrast, the Meta Quest 3 exhibited poor calibration performance, making it unsuitable for accurate control of colors. Moreover, we showed that the color reproduction accuracy depends greatly on the region of the lens. The sweet spot area of the lenses consistently showed way better results than the peripheral areas. These variations underscore the importance of device-specific considerations in the development and application of psychophysical experiments using VR.

## 5 Conclusion

This study introduced a new approach to calibrate VR HMDs, aiming to resolve some issues present in previous methods. We showed that it is possible to perform successful color calibration and accurately control stimulus colors without the drawbacks of previous solutions. Indeed, our approach can disable tone mapping only on specific objects in the scene, allowing for a globally realistic scene while accurately controlling the physical color of a stimulus. Our approach can also be used to disable tone mapping for the entire scene globally instead of at the specific object level.

In this sense, our approach is very flexible and adaptable to various scenarios needed by researchers. We also explained that another advantage of our solution is the use of standard Lambertian materials, which have normal behavior with respect to lighting, unlike the unlit materials used in previous studies.

Moreover, we developed our solution using Three.js instead of the commonly used Unity or Unreal Engine. To our knowledge, this methodology is new in vision science studies and is intrinsically interesting by bringing more diversity and options to researchers. In addition, this 3D engine offers other advantages, such as transparency, portability, and ease of use. We also provide source code to ensure easy reproduction of our method.

Finally, we showed a comparison of the calibration performances for three different HMDs. We highlighted that the HTC Vive Pro Eye performs excellently, the Meta Quest Pro performs well, and the Meta Quest 3 performs poorly. We also showed that the color reproduction fidelity varies significantly across the lenses. It is optimal where the user’s eye is positioned by default, but it drops in the peripheral areas. This is especially the case for the Meta Quest Pro and Meta Quest 3. Therefore, we warned that even if it is possible to precisely control the color produced in the center of the lenses, it is much more challenging to control it in the peripheral regions. This is something to take into account when designing psychophysical experiments. Indeed, it is unlikely that an observer would look at the scene or a stimulus only through the center of the lens.

In conclusion, this study offers a framework and insightful information for the design and utilization of VR in vision research. The ability to calibrate and control color accurately in VR HMDs without other drawbacks could significantly enhance research by providing new ways to design and run psychophysical experiments. Future research should explore the integration of this method into real-world applications, such as VR psychophysical experiments where color accuracy is critical.

## 6 Acknowledgments

This work was supported by JST SPRING, Japan Grant Number JPMJSP2106 to KD, and JSPS KAKENHI Grant Number 23K28174 and 21KK0203 to TN.

## Commercial relationships

none

## References

Brainard, D. H. (1989). Calibration of a computer controlled color monitor. Color Research & Application, 14(1), 23–34.

Brainard, D. H., Pelli, D. G., & Robson, T. (2002). Display characterization. Signal Process, 80, 2–067.

Díaz-Barrancas, F., Gil-Rodríguez, R., Aizenman, A., Bayer, F., & Gegenfurtner, K. (2023). Color calibration in virtual reality for unity and unreal. In 2023 IEEE Conference on Virtual Reality and 3D User Interfaces Abstracts and Workshops (VRW) (pp. 733–734). IEEE.

Gil Rodríguez, R., Bayer, F., Toscani, M., Guarnera, D., Guarnera, G. C., & Gegenfurtner, K. R. (2022). Colour calibration of a head mounted display for colour vision research using virtual reality. SN Computer Science, 3, 1–10.

Gil Rodríguez, R., Hedjar, L., Toscani, M., Guarnera, D., Guarnera, G. C., & Gegenfurtner, K. R. (2024). Color constancy mechanisms in virtual reality environments. Journal of Vision, 24(5), 6–6. Retrieved from doi: 10.1167/jov.24.5.6

Koida, K. (2021). About color calibration of displays. Retrieved 2023-12-07, from https://www.eiiris.tut.ac.jp/koida/colorcalib/

Motoyoshi, I., & Matoba, H. (2012). Variability in constancy of the perceived surface reflectance across different illumination statistics. Vision Research, 53(1), 30–39.

Sharma, G., & Bala, R. (2017). Digital color imaging handbook. CRC press.

Toscani, M., Gil, R., Guarnera, D., Guarnera, G., Kalouaz, A., & Gegenfurtner, K. R. (2019). Assessment of OLED head mounted display for vision research with virtual reality. In 2019 15th International Conference on Signal-Image Technology & Internet-Based Systems (SITIS) (pp. 738–745). IEEE.

Wilson, A., & Hua, H. (2021). Design of a pupil-matched occlusion-capable optical see-through wearable display. IEEE Transactions on Visualization and Computer Graphics, 28(12), 4113–4126.

Zaman, N., Sarker, P., & Tavakkoli, A. (2023). Calibration of head mounted displays for vision research with virtual reality. Journal of Vision, 23(6), 7–7. Retrieved from doi: 10.1167/jov.23.6.7

